# Body size shifts influence effects of increasing temperatures on ectotherm metabolism

**DOI:** 10.1101/139279

**Authors:** Kristina Riemer, Kristina J. Anderson-Teixeira, Felisa A. Smith, David J. Harris, S.K. Morgan Ernest

## INTRODUCTION

Environmental temperature influences organisms in many ways; temperature increases or decreases rates of physiological processes (Brown et al., 2012), determines timing of reproduction (Olive, 1995), and even directly affects mortality (Pauly, 1980). Because of the far-reaching influence of temperature, projected increases in global temperatures due to climate change are expected to substantially alter diverse species characteristics. Increased temperatures have already been implicated in shifts in species geographic distributions (e.g., Buckley et al., 2010), and in the phenology of species’ life history and development (e.g., Wolkovich et al., 2012).

It is predicted that global warming will also increase metabolic rates of ectotherms (Seebacher et al., 2015; Dillon et al., 2010). Metabolic rate is a key physiological process that represents the rate of energy required for the maintenance, growth, and reproduction of organisms. Temperature influences metabolic rates in ectotherms through its influence on the kinetic energy available for chemical reaction. Because the relationship between temperature and metabolic rate is positive and exponential until an upper temperature threshold, small changes in temperature can have substantial impacts on metabolic rate (Gillooly et al., 2001). By directly increasing ectotherm metabolic rates, warmer temperatures would have considerable impacts on the ecology of communities and ecosystems (Anderson-Teixeira et al., 2012; Lemoine & Burkepile, 2012; O'Connor et al., 2011; Rall et al., 2010). Changes in metabolic rate affect many aspects of organismal biology and ecology, including individual fitness (Burton et al., 2011), population dynamics (Buckley et al., 2014), community composition (Marquet et al., 2004), and even ecosystem processes such as ecosystem respiration and nutrient cycling (Gilbert et al., 2014; McIntyre et al., 2008). Therefore, understanding the impact of temperature increases on metabolic rates is critical to determining how ectotherm organisms, and the ecological communities in which they perform key roles, will response to climate change.

While the direct effect of temperature on ectotherm metabolic rates is well-known, temperature also affects other aspects of organismal biology which in turn influence metabolic rate. One such organismal characteristic that influences metabolic rate and is influenced by temperature is body size. Temperature affects body size, a phenomenon often referred to as the size-temperature rule in which ectothermic individuals commonly develop to a smaller adult size when raised at higher temperatures (Forster et al., 2011a; Angilletta, 2004; Atkinson, 1994). This shift in adult size occurs quickly, with individual body size often responding to temperature change after a single generation (e.g., fruit flies in Partridge et al., 1994). Smaller ectotherms then have decreased metabolic rate due to the positive allometric relationship between body size and metabolic rate (Kleiber, 1932; West et al., 1997; Brown et al., 2004). Thus, in addition to its direct effect on metabolic rate, temperature influences metabolic rate indirectly via body size shifts.

Predictions of ectothermic metabolic response to warming temperatures have focused on the direct effect of temperature on metabolic rate while neglecting the indirect effects from size shifts (Dillon et al., 2010). This occurs even though the relationships between temperature and metabolic rate, size and metabolic rate (Gillooly et al., 2001; Brown et al., 2004), and temperature and size (Walters & Hassall, 2006) are well documented. Indirect effects often emerge in ecology because ecological systems are complex systems composed of multiple, and sometimes antagonistic, interactions. These indirect effects can be strong enough to modify outcomes, rendering predictions that only consider direct interactions incorrect (e.g., apparent competition in Holt, 1977). While warmer temperatures from climate change will directly result in increased metabolic rates, this increase could be offset by a decrease in metabolic rates due to the concurrent indirect effect of smaller body size.

Here we examine how predictions of ectotherm metabolic rates differ when the indirect effect of increased global temperatures on size is included in addition to the direct effect. While higher temperatures cause metabolic rates to increase when size is constant (i.e., direct effect), this increase could be dampened when size is allowed to vary (i.e., indirect effect) consistent with empirical data, if not result in an overall decreased metabolic rate. To estimate how much size changes in response to temperature, we collect experimental data on observed ectothermic size shifts in response to temperature changes. We then use a previously established model that integrates size and temperature effects on metabolic rate to estimate metabolic rates when size is constant and when size varies. We compare these metabolic rates to determine how this established indirect effect of increased temperature alters the magnitude and direction of existing predictions that use only the direct effect.

## METHODS

### Data

We used data from published experimental studies that raised individuals of ectotherm species at constant temperatures. From these studies, we obtained average adult size of all individuals grown at each temperature treatment. Criteria for inclusion were that (1) in each study individuals were raised at a minimum of two experimental temperatures, (2) individuals were either lab-bred or collected at an early life stage, and (3) sufficient food was provided so that resource limitation did not influence ontogenetic growth. These criteria were laid out by Forster et al. (2011a), which was also the source of most of our data, and we collected some additional data from the literature that also conformed to these criteria (Coker, 1993; Berven, 1982; Stacey & Fellowes, 2002; Marti & Carpenter, 2008; Oetken et al., 2009). Some studies had multiple trials to compare responses of individuals from different latitudes or elevations; we retained data for all trials and they were kept separate for the analysis. If length was the only size metric provided, we converted it to mass using allometric relationships (Appendix S1 in Supporting Information).

Studies examined body size response to temperature across a range of temperatures (2°C – 36°C) that differed from each other by various temperature increments (1°C – 29°C difference in temperature between experiments within studies). To simplify this analysis and focus on our core question of whether the indirect temperature effect on body size could, on average, offset or substantively ameliorate the direct effect of temperature on metabolic rate, we further filtered this data by only considering pairs of experiments within a study that differed by 3°C in experimental temperature. We chose 3°C as our temperature difference because it is within the bounds of predicted future temperature change from climate models, though particular species may or may not experience this specific temperature increase in their native ranges. From each 3°C experimental pair, we considered the mass value associated with the lower temperature to represent that species' mass before temperature increase while mass value reported for the higher temperature represented size response to a 3°C increase in temperature. Some studies had multiple experimental pairs whose experimental temperatures differed by 3°C (i.e., one pair of 15°C and 18°C, and a second pair of 18°C and 21°C); in these cases we kept all 3°C pairs. This filtered subset of the data had temperature ranges of 3°C to 30°C for the lower temperature and 6°C to 33°C for the higher temperature.

The final dataset contained 191 pairs of average adult masses for 45 species across 40 studies. This dataset includes species from seven taxonomic classes, ranging from Insecta to Amphibia, and terrestrial species from every continent except Antarctica and aquatic species from every ocean and many large bodies of water. Most species are very small (<100 mg) invertebrates because available data for larger ectotherms was limited. Mass values span five orders of magnitude, from 2 µg to 100 mg. Data and code have been deposited in the online Dryad Data Repository (http://datadryad.org).

### Metabolic rates

Size-temperature studies do not typically measure metabolic rate. Therefore, we used a model to calculate species’ metabolic rates from the experimental temperature-mass dataset. This model is central to the metabolic theory of ecology (MTE) (Brown et al., 2004) and combines both mass and temperature effects on metabolic rates:

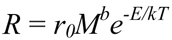

where *R* = metabolic rate, *r_0_* = scaling constant, *M* = mass (g), *b* = unitless scaling exponent, *E* = activation energy of respiration (eV), *k* = Boltzmann's constant (8.617 × 10^-5^ eV/K), and *T* = temperature (K). Though the mechanism underlying the MTE equation has been questioned (O'Connor et al., 2007), it provides a reasonable empirical approximation across a wide range of taxa (Gillooly et al., 2001) and has been used previously to estimate metabolic rates response to temperature change (e.g., Dillon et al., 2010). An advantage of this equation for our study is that it allows us to incorporate both the direct effect of temperature on metabolic rate (*e^−E/kT^*) and the indirect effect through organismal size (*M^b^*).

We used values of *b* and *E* specific to each taxonomic class to allow for taxon-specific responses of metabolism to temperature. Because availability of class-specific *b* and *E* values varied widely among the classes, we used different approaches for determining these values. When data on organismal metabolic rate available for a class also contained size and temperature information (Branchiopoda, Amphibia, Malacostraca, Maxillopoda; Makarieva et al., 2008; White et al., 2012), we used a multiple regression method (White et al., 2012) to calculate *b* and *E*. Values for Actinoperygii were estimated using fish species from several classes (Gillooly et al., 2001) and values for Insecta were acquired from literature sources (Chown et al., 2007; Irlich et al., 2009), while the average values of *b* = 0.75 and *E* = 0.63 eV (Brown et al., 2004) were used for Entognatha, the most data-limited class. Data used to calculate *b* and *E* were non-overlapping with our experimental temperature-mass dataset.

For each species, we used the MTE equation to calculate how much metabolic rate changed due to only the direct effect of temperature increase, and from both the direct effect of temperature and indirect effect of the empirical body size response to temperature. To do so, we calculated three metabolic rates for each 3°C experimental pair in the temperature-size dataset (Fig. 1A):

i. “Starting metabolic rate” represents the metabolic rate prior to temperature increase, and was calculated using size and temperature data from the lower temperature experiment of each pair.
ii. “Constant size metabolic rate” is the hypothetical metabolic rate including only the direct effect of temperature increase. It was calculated with the size from the lower temperature of each experimental pair, and temperature from the higher temperature of each pair (i.e., the assumption that size at the lower temperature does not change as temperature increases).
iii. “Varying size metabolic rate” represents the empirical metabolic rate that includes both the direct and indirect effects of temperature increase. It was calculated using both size and temperature data from the higher temperature of each pair.

**Figure 1.**
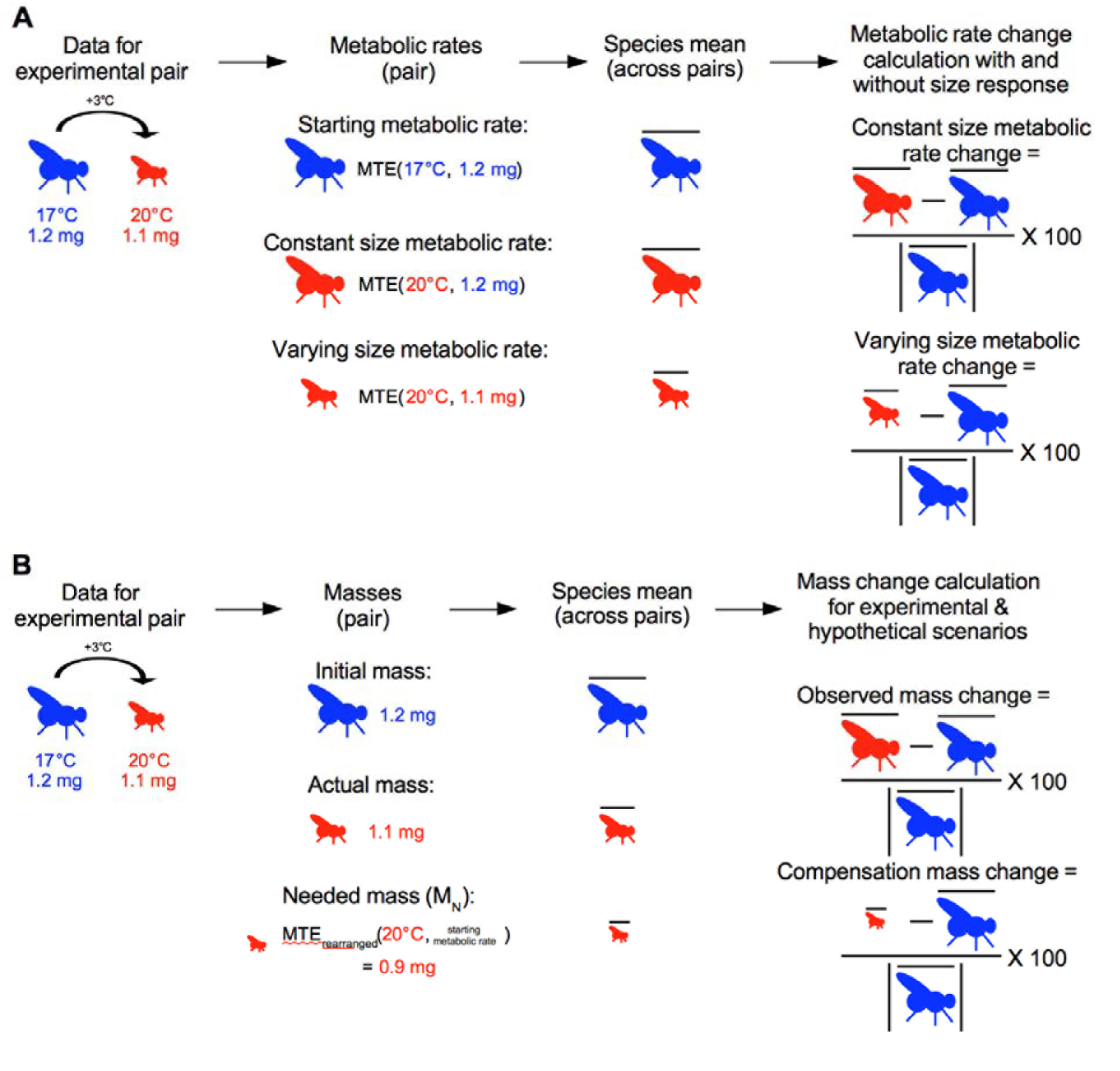
Diagrams showing how the experimental data from the pairs were used to calculate constant size and varying size metabolic rate changes for each species (**A**), and how the data were used to calculate observed and compensation mass change for each species (**B**). The example data is from the species *Aedes aegypti*.

#### Comparison

To visualize the difference between the metabolic predictions including only the direct effect of temperature versus including both the direct and indirect effects of temperature, we examined how much varying size metabolic rate increased from initial compared to constant size metabolic rate. Because some species had data from multiple 3°C experimental pairs, we condensed the data by calculating species averages for starting metabolic rate, constant metabolic rate, and varying size metabolic rate (Fig. 1A). We used these average values to calculate i) “constant size metabolic rate change”: the percent change from starting metabolic rate to constant size metabolic rate and ii) “varying size metabolic rate change”: the percent change from starting metabolic rate to varying size metabolic rate. These two change metrics represent how much metabolic rate changes due to only the direct effect of temperature increase (i.e., constant size metabolic rate change), and when the indirect effect of temperature, in addition to the direct effect, is included (i.e., varying size metabolic rate change). Positive percent change showed an increase in metabolic rate from starting metabolic rate for each species. To assess the difference between metabolic rates with and without the indirect effect, each species’ constant size and varying size metabolic rates were log-transformed and compared across species with a paired t-test.

#### Linear mixed model

We used a mixed model to determine if other factors, besides the body size response, impacted the difference between constant size and varying size metabolic rates. The response variable for the model was the log of the ratio between each pair's constant size and varying size metabolic rates, where a positive log-ratio indicated that the pair's varying size metabolic rate was smaller than its constant size metabolic rate. In addition to the model results, likelihood ratio tests were used to determine each effect's significance. We included absolute temperature, as represented by the lower temperature for each pair, as a fixed effect because higher starting temperatures are expected to have a disproportionate effect on metabolic rates due to their exponential relationship. The random effects in the model were the taxonomic classifications of species and class, as metabolic rate varies amongst these groupings due to biology and ecology. We initially included study and trial as random effects, to take into account differences in experimental setups, but these were not included in the final model because their maximum likelihood estimates for variance were near zero and they therefore had no impact on the final model. Similarly, because metabolic rate depends on the relative size of organisms, we included mass from the lower temperature of each pair as a fixed effect but it was removed because it did not have a substantial effect on the metabolic rate log-ratio (χ^2^ = 0.7015; df = 1; p = 0.4). We ran the linear mixed model using the R package lme4 version 1.1.9 (Bates et al., 2015; Winter, 2013).

### Compensation mass

We assessed the magnitude of size response to temperature needed to offset the direct effect of temperature on metabolic rate, and how close each species' observed size response to temperature came to reaching this predicted value (Fig. 1B). To calculate the predicted size necessary to offset the 3°C increase in temperature for an experimental pair (MN), we rearranged the MTE equation to solve for size:

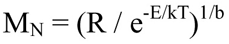

where R is the starting metabolic rate and T is the observed temperature from the higher temperature experiment for each pair. Thus, needed mass (M_N_) is the mass a species would have to be under higher temperature conditions in order for metabolic rate to not change.

We calculated how much each species' size actually changed with increased temperature, (“observed mass change” = percent change from initial mass to actual mass), and how much size theoretically needed to change to maintain metabolic rate with increased temperature (“compensation mass change” = percent change from initial mass to needed mass) (Fig. 1B). Observed mass changes or compensation mass changes of less than 100% indicated that the mass of a species either did or needed to decrease, respectively. A paired t-test was used to compare log-normalized actual and needed mass values. All analyses were completed using R version 3.3.1 (R Core Team, 2016).

## RESULTS

### Metabolic rates comparison

Metabolic rates differed when the indirect effect of temperature increase was included with the direct effect. All varying size metabolic rate changes, which included the indirect effect of temperature on size, were positive (average percent change = 23%; Fig. 2), indicating that ectotherm metabolic rate will still increase with temperature despite decreases in body size. However, most species' metabolic rates did not increase as much when the indirect effect was included, as shown by smaller varying size metabolic rate changes than constant size metabolic rate changes (Fig. 2). There was also a statistically significant difference between these metabolic rate changes, with varying size metabolic rate being consistently smaller than constant size (t_44_ = 3.34; p = 0.002; 95% CI = 0.017 – 0.070). While the majority of species had a smaller increase in metabolic rate with the body size response included, for 9 of the 45 species incorporating size shifts actually caused metabolic rate to increase even more than expected from just the direct effect of temperature.

**Figure 2.**
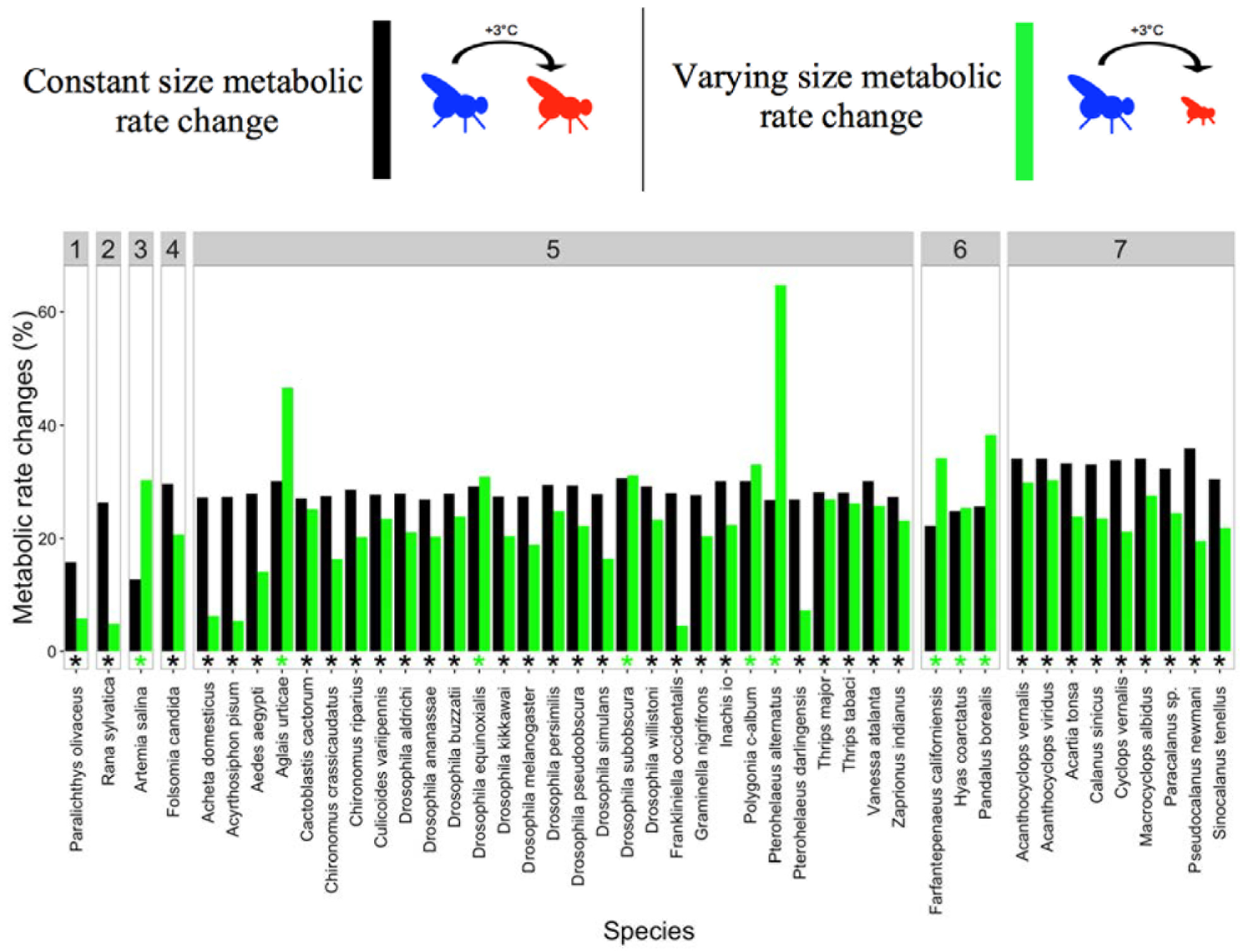
Constant size metabolic rate change (black bar) and varying size metabolic rate change (green bar) for all species, represented as percent change from starting metabolic rate to metabolic rate without the indirect effect of size response and to metabolic rate with indirect effect, respectively. A positive percent difference indicates an increase from starting metabolic rate. Asterisks are colored according to whether constant size metabolic rate (black asterisk) or varying size metabolic rate (green asterisk) is larger for each species. Species are grouped by class, with classes labelled at the top of the figure as follows: 1 = Actinoperygii, 2 = Amphibia, 3 = Branchiopoda, 4 = Entognatha, 5 = Insecta, 6 = Malacostraca, 7 = Maxillopoda.

### Linear mixed model

Similarly, most pairs had smaller varying size metabolic rates than constant size metabolic rates. Of 191 pairs, 82% had positive log-ratios, which represent the difference between constant size and varying size metabolic rates. These log-ratios were not explained by the three factors of interest we included in the model. While absolute temperature (χ^2^ = 9.27; df = 1; p = 0.002), taxonomic class (χ^2^ = 3.90; df = 1; p = 0.048), and taxonomic species (χ^2^ = 5.48; df = 1; p = 0.019) did influence the differences between metabolic rates, they were not sufficiently biologically significant. With increasing absolute temperature, varying size metabolic rate tended to get increasingly smaller than constant size metabolic rate, as shown by a slightly positive slope in the relationship between absolute temperature and log-ratio (slope = 0.004 ± 0.001).

However, this increasing difference had minimal biological relevance because it was between one and two orders of magnitude smaller than the actual difference between metabolic rates. Similarly, species classification explained only 17% of the variability in random effects, while class explained a more substantial 43% of random effects variability (Fig. 2).

### Compensation mass

No species' mass decreased enough for their metabolic rate to remain constant regardless of temperature increase. To retain constant metabolic rates, all species would need to get smaller (Fig. 3). Actual mass (i.e., empirical mass in response to temperature increase) and needed mass (i.e., theoretical mass required in order for metabolic rate to not change due to temperature increase) were statistically significantly different, with needed masses smaller than actual masses (t_44_ = 17.707; p < 2.2×10^-16^; CI = 0.25 – 0.32). While most species empirically decreased in size, 20% of species actually increased in size in response to temperature increase (observed mass change mean ± standard deviation: -4% ± 11%). These were the same species that had a greater increase in metabolic rate when the body size response was included (Fig. 2).

**Figure 3.**
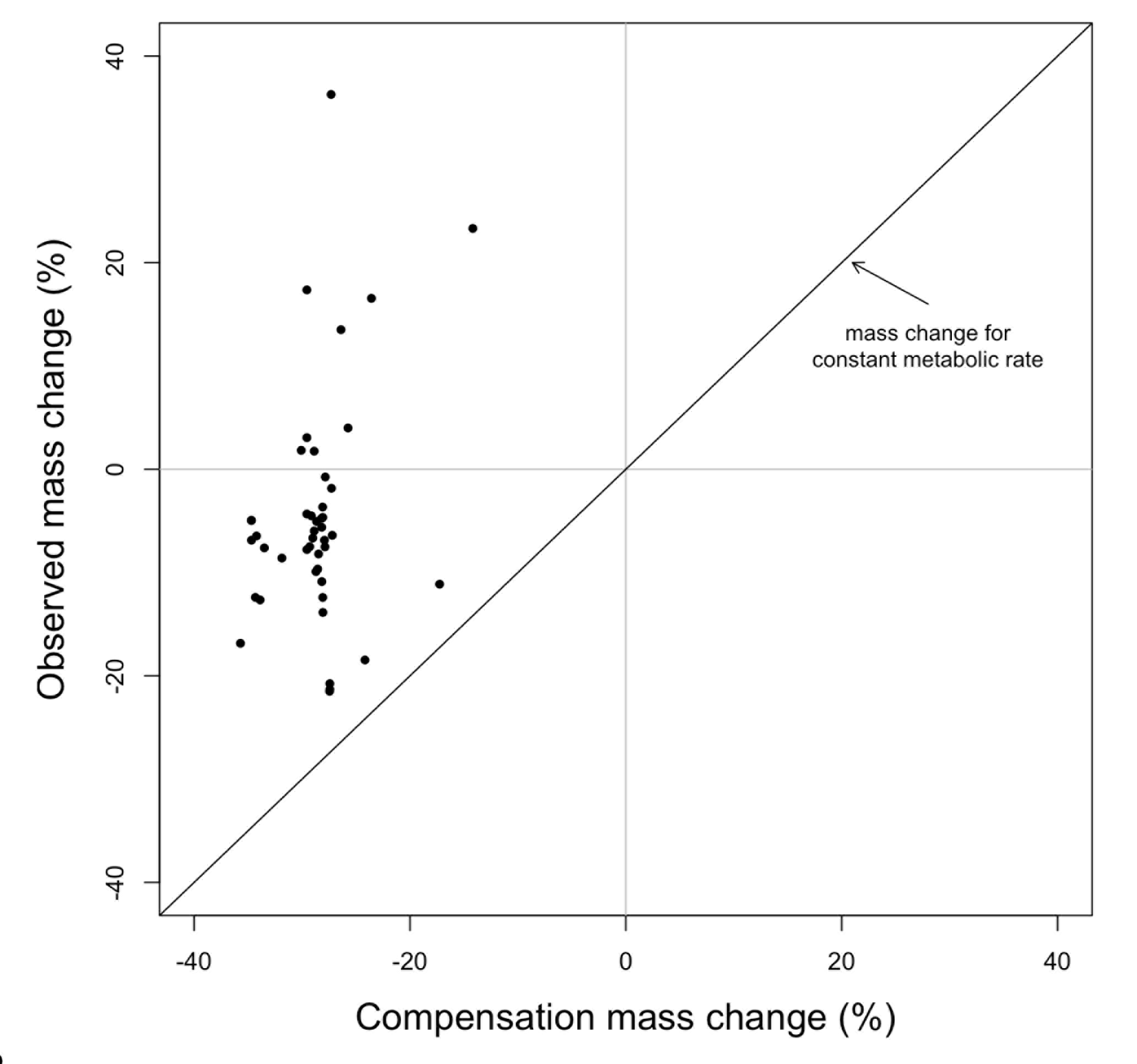
Comparison between compensation mass change and observed mass change for each species. Compensation mass change is the percent change from initial mass to needed mass (i.e., mass change needed to maintain metabolic rate with increased temperature), while observed mass change is the percent change from initial mass to actual mass (i.e., observed mass change of individuals of each species raised experimentally). No change in mass and no change needed in mass are indicated by grey lines. Equivalence between needed mass and actual mass shown by black line.

### DISCUSSION

Consistent with expectations, predictions of most species' metabolic rates were smaller when the indirect effect of temperature was included with the direct effect. While all metabolic rates increased with increased temperatures whether or not the indirect effect was included, incorporating the body size response to temperature significantly dampened the increase in metabolic rate for most species (Fig. 2). Species decreased in size by up to 20% (Fig. 3). Though there is no accepted quantitative description of the relationship between temperature and size for ectotherms (Forster et al., 2011b), the magnitude of size decrease by these species was similar to results from previous studies. For example, worm species *Caenorhabditis elegans* was 6% smaller when raised at a temperature increase of 5°C (Voorhies, 1996) and two mayfly species *Ameletus ludens* and *Ephemerella subvaria* were 6% and 26% smaller, respectively, at a 3.1°C warmer temperature (Sweeney & Vannote, 1978). None of the species in this study decreased sufficiently in size for the indirect effect of temperature to offset the direct effect. All species would have needed to decrease between 15% and 35% in size for their metabolic rates to remain constant with temperature increase (Fig. 3).

A small proportion of species increased in size, instead of decreasing as expected, in response to increased temperature. Of 36 species, 9 species increased in size (Fig. 3) and therefore had a greater increase in metabolic rate when the indirect effect of temperature was included (Fig. 2). These included four crustacean species from two classes (*Artemia salina, Farfantepenaeus californiensis, Hyas coarctus, Pandalus borealis*) and insect species including two butterfly species (*Aglais urticae, Polygonia c-album*), two out of 11 fruit fly species (*Drosophila equinoxialis, Drosophila subobscura*), and one crop pest (*Ptero alternus*). This opposing size response is consistent with previous research as, for example, 12% of the studies in the meta-analysis by Atkinson (1994) showed a positive relationship between temperature and size. While it is unknown why some species exhibit this opposing response, many explanations have been proposed based on growth rates (Angilletta & Dunham, 2003; Walters & Hassall, 2006), abiotic conditions (Atkinson, 1994), extreme temperatures (Kingsolver & Huey, 2008), and life history characteristics (Walters & Hassall, 2006; Forster et al., 2012). Regardless of the mechanism, these anomalous species will have relatively greater metabolic rates and body sizes from increased temperatures, with the accompanying higher energy requirements. Their greater use of space, food, and other resources could result in disproportionately greater ecological impacts and ecological mismatches, such as substantial changes in prey abundances due to increased consumption by predators (Rall et al., 2010).

Though there was a significant difference in species metabolic rates when the indirect effect of temperature was included, the biological relevance of this difference is unknown. Including the indirect effect of temperature on size generally resulted in small changes in the predicted metabolic rate. Species that declined in size with increasing temperature experienced, on average, a 9% reduction in the predicted metabolic rate increase, while those that increased in size saw an average 11% increase in metabolic rate from that expected due to the direct effect of temperature alone. This magnitude of difference in metabolic rate seems relatively small, but its ecological relevance could depend on context. For example, Gilbert et al. (2014) showed that small changes in biomass potential resulted in large and unpredictable fluctuations in food web stability but only when biomass potential was low. Thus, whether or not these small changes in expected metabolic rates due to the size response to increasing temperature will have cascading effects on the ecology of those species or the communities they inhabit may depend on the productivity of the system and their trophic interactions. Further work would be needed to determine if there is an ecological impact due to the difference in metabolic rate caused by the body size response.

Because none of the species in this study decreased enough in size to offset the expected direct effect of temperature on metabolic rate, increasing metabolic rates is still likely to be a widespread response of ectotherms to warming. The overall increase in ectotherm metabolic rates will have substantial consequences for every aspect of ecological systems, from population-level and up (Schmidt-Nielsen, 1984; Anderson-Teixeira et al., 2012). Individuals of each species will require more energy to maintain normal maintenance, growth, and reproductive function and therefore use a greater proportion of resources. Assuming resource availability remains the same, this will increase intraspecific competition, resulting in lower species abundances and higher extinction probabilities (MacArthur & Wilson, 1963; Hurlbert & Stegen, 2014) that will shift community composition, potentially changing the dominant species, trophic structure, and interspecific interactions. This would ultimately affect ecosystem-level energy fluxes (Allen et al., 2005; Yvon-Durocher & Allen, 2012). Like metabolic rate, size itself impacts many aspects of population, community, and ecosystem ecology (e.g., cascading effects on foraging and interspecific competition in woodrats in Smith et al., 1998) independently of its impact through metabolic rate. The shifts in size that ectotherms may experience with warming are likely to compound many of these cascading impacts as well. Incorporating the effects of temperature change through its coordinated impacts on metabolism and size will be challenging, but previous work on changing body size (Woodward et al., 2005; Ohlberger, 2013) is a potential guide. Particularly informative is an extensive study by O'Gorman et al. (2012) which showed significant effects of body size on individuals, populations, communities, and ecosystem processes in Icelandic streams with a temperature gradient.

Understanding how climate change will impact species and ecosystems is difficult because ecological systems are complex. Temperature does not impact one aspect of ecology, but influences dynamics at almost every level, from the physiology of organisms to the rates of ecosystem fluxes. We often reduce this complexity by studying the direct effects of temperature change on some particular component of ecology, such as the effect on metabolic rate, and then sometimes additionally the direct effect of the changed ecological component on some other ecological process. Our results highlight the need to consider the coordinated impact of multiple pathways of effects when making predictions about how species will respond to climate change. While we focused here on the body size response of ectotherms to temperature as an indirect effect on metabolic rate, this is by no means the only indirect pathway by which temperature can impact metabolic rate. Temperature can impact size through a variety of other environmental pathways besides the temperature-size response, including through resource availability, which could then consequently affect body size in various ways (McNab, 2010; Bickford et al., 2011). Designing studies to assess the relative importance of direct and indirect effects of temperature on species, communities, and ecosystem properties of interest is an important next step for our ability to predict the impacts of climate change.

## ACKNOWLEDGEMENTS

This research was supported by the Gordon and Betty Moore Foundation Data-Driven Discovery Grant and the NSF CAREER grant, both to Ethan White. Thanks to Ellen Bledsoe for feedback on data visualization.

## SUPPLEMENTARY MATERIALS LIST

**Appendix S1** Methods for converting non-mass measurements to mass

## DATA ACCESSIBILITY

All temperature-mass data, along with associated cleaning and analysis R scripts, are available on Dryad.

## BIOSKETCH

The Weecology group, in conjunction with collaborators, works on diverse ecological questions by linking empirical and quantitative approaches, with a focus on macroecology and computational techniques.

Author contributions: K.A.T., F.S., and S.K.M.E conceived the project and collected data, K.R., D.J.H., and S.K.M.E. analyzed data, and K.R. and S.K.M.E. primarily completed the writing with contributions from all co-authors.

